# DnaK refolds denatured proteins by actively pulling out their misfolded structural elements

**DOI:** 10.1101/2025.09.22.677870

**Authors:** Oskar K. Marszałek, Piotr E. Marszalek

## Abstract

DnaK, a prokaryotic Hsp70 chaperone, plays a central role in proteostasis by restoring native structures to heat-denatured proteins in an ATP-hydrolysis–dependent manner. While structures of DnaK in complex with nucleotides, co-chaperones, and short peptides have been resolved, structures with larger, stably folded substrates—such as firefly luciferase (Fluc, 61 kDa)—are lacking, limiting mechanistic understanding of how DnaK refolds such proteins.

Here, we generated models of the DnaK–Fluc complex using AlphaFold3 and evaluated their mechanistic relevance. In one of three major model clusters, Fluc is unexpectedly immobilized beneath the DnaK α-helical lid against the nucleotide-binding domain (NBD), rather than interacting primarily with the substrate-binding domain β (SBDβ), as commonly assumed. All-atom molecular dynamics simulations indicate that, in this configuration, the lid can engage a thermally destabilized Fluc helix (residues 405–411), which we recently identified as the first—and likely the only—helix to irreversibly melt at 42 °C.

Upon binding, the lid forms extensive hydrogen-bonding interactions with the melted helix. These interactions persist during lid movement toward SBDβ (following ATP hydrolysis), enabling the lid to actively extract the helix from the Fluc surface. In contrast, simulations with the helix in its native folded state show that the lid cannot extract it, leaving the native structure unaffected. Equilibrium simulations further indicate that, once extracted and mechanically stretched, the melted helix can refold to its native conformation.

Together, these findings suggest a revised mechanism for DnaK-mediated protein refolding, in which the α-helical lid selectively recognizes structurally compromised segments, forms stabilizing hydrogen bonds, and—powered by ATP hydrolysis—mechanically pulls them away from the protein surface to facilitate their refolding.

**SIGNIFICANCE:** DnaK is a model chaperone, which can reactivate thermally denatured proteins. Over the span of 40 years, significant findings have been made about DnaK’s structure, dynamics and interactions with its co-chaperones, the exact molecular mechanism by which DnaK refolds misfolded proteins remains a mystery. This work exploited Alphafold3 to generate atomistic models of complexes between DnaK and Firefly luciferase. Molecular dynamics simulations directly captured how DnaK may assist thermally denatured proteins by mechanically pulling out their misfolded helices. This study provides a new insight into the DnaK mechanism.

## 1. INTRODUCTION

DnaK is an *E. coli* Hsp70 chaperone (70 kDa heat shock protein) with many cellular functions. It is involved in transporting proteins across membranes, assisting nascent polypeptide chains to acquire their functional structures, refolding polypeptides denatured by thermal shock, and more generally in inhibiting protein aggregation and maintaining protein homeostasis through cooperation with other chaperones (1–7). The DnaK polypeptide chain has 638 amino acids, and it folds into two distinct domains: the N-terminal, nucleotide binding domain (NBD) and the C-terminal substrate binding domain (SBD) (Fig. 1). Both domains have internal subdomains whose movement is strictly coordinated by and coupled with other molecules: substrate and co-chaperone binding (DnaJ and GrpE (8)), ATP hydrolysis and controlled ADP release (9,10). The SBD is composed of two mobile parts. The first part is the cradle-like SBDβ, which accommodates short unstructured peptides, such as a model NR peptide (11) or unstructured segments of larger proteins, such as σ^32^ (12) or proPhoA (13). The second part, SBDα, consists of a long helical lid structure, with a characteristic triple-helical bundle. It, in turn, is followed by a C-terminal unstructured segment whose function is unknown (14).

**Figure 1.**
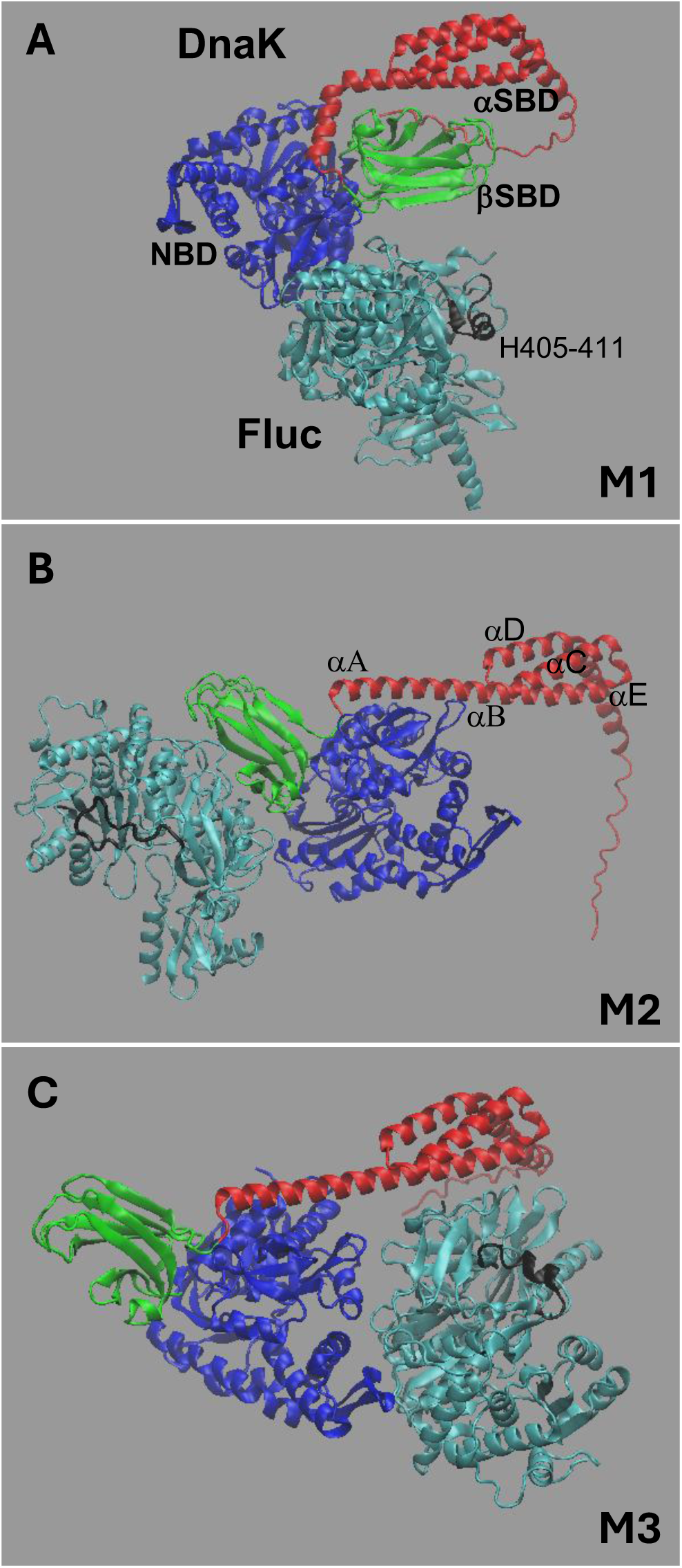
Highest scoring representatives of DnaK-Fluc models generated by AF3. All models were grouped into 3 clusters (M1-like, M2-like, and M3-like) that had common configurations of DnaK and Fluc.

DnaK is an exquisite nanomechanical machine with sensors and actuators that can produce and consume chemical energy to perform multiple functions (1,15). When ATP is bound within the NBD, DnaK is configured in the so-called open conformation where the SBDα lid is separated from the SBDβ and attached to the NBD (14,16). After ATP hydrolysis, with ADP still present in the NBD, the lid moves back to its position over the SBDβ, which becomes slightly separated from the NBD, creating the so-called closed conformation of DnaK (17). The lid moves back toward the NBD after ADP is removed from the NBD with help from the GrpE co-chaperone (18,19). Although there is an essential understanding supported by strong evidence through many papers (e.g. (12,20) of both DnaK’s structural details and the allosteric communication between its NBD and SBD, it remains less clear how these observations can be combined and applied to understand the mechanism by which DnaK assists thermally denatured proteins during the heat shock response. Several hypotheses were proposed over the years, including the holdase model and (un)foldase models. The latter proposes two variants, one involving so-called entropic pulling (21) and another involving active mechanical substrate unfolding by DnaK, using energy from ATP hydrolysis, for some unspecified mechanical action (22–26). All these conjectures are partially supported by experiment and modeling efforts that are summarized in several reviews (e.g. (1,27,28)).

In our opinion, the difficulties in gaining a better understanding of how DnaK refolds denatured proteins are related, in part, to the absence of structural information on (transient) complexes between DnaK and its larger substrates, such as firefly luciferase (Fluc), which has been the gold standard in research on chaperone mechanisms (29,30). Additionally, structural alterations triggered at heat shock temperatures (e.g. 42 °C) in DnaK substrates, such as those in Fluc, are also not documented, further impeding conceptualization of possible DnaK mechanisms to eliminate these structural defects.

Recently, we addressed the second of the above-mentioned difficulties by carrying out an extensive computational study of structural alterations that may occur at heat shock temperatures in Fluc (31). Surprisingly, multiple very long molecular dynamics simulations of Fluc at 42 °C revealed relatively small but *irreversible* misfolding events involving just a single 2-turn helix, encompassing residues 405 to 411 (H405-411) (31). The thermal melting of this helix likely affects Fluc’s catalytic bioluminescence activity, which is used as a proxy for the structural integrity of Fluc in chaperone-mechanism studies. This observation prompted us to think about how DnaK could act to refold this misfolded helix.

Because no experimental structures—even partial ones—exist for DnaK–Fluc complexes, we used Alphafold3 (AF3) (32) to generate structural models of such a complex. Exploring a large combination of user-provided input parameters, AF3 predicted three distinct classes (clusters) of complexes between DnaK and Fluc (Fig. 1). In two of these clusters, including the highest scoring models, Fluc is surprisingly, not associated with SBDβ of DnaK, as typically assumed, but is associated with either the NBD alone (cluster one, Fig. 1*A*) or is placed under the alpha helical lid and associated with the NBD on the side opposite to SBDβ (cluster 3, Fig. 1*C*). In cluster 2, Fluc is associated with SBDβ (Fig.1*B*), as anticipated from the structural studies of DnaK complexes with unfolded peptides (17). Here, we analyze these AF3 models of DnaK-Fluc complexes and employ molecular dynamics simulations to examine how DnaK may interact with misfolded Fluc to assist its refolding. Although speculative at present, our observations suggest a possible new DnaK mechanism for refolding of misfolded proteins, in which the DnaK alpha helical lid engages misfolded structural elements through hydrogen bonding and uses the energy from ATP hydrolysis to forcefully pull them out from the protein surface, so they can recover their native fold upon relaxation.

## 2. MATERIALS AND METHODS

### 2.1 Generating models of DnaK-Fluc and DnaK-Fluc-DnaJ complexes

AF3 jobs were run by specifying target proteins sequences, UniProt ID: P0A6Z0 for DnaK, P08659 for Fluc and P08622 for DnaJ. Using the WEB-based AF3 interface we specified modeling conditions: such as instructions for using or not internal AF3 PDB templates, including or not ligands (ATP) and ions (Na, Cl). For local jobs (Duke University Computing Cluster, DCC) we also could use various combinations of MSA information (unpaired MSA, paired MSA, both, none) and could supply custom protein templates. As a custom DnaK template we used PDB: 4B9Q (14). As a custom Fluc template, we used our own structure generated through molecular dynamics simulations of 1BA3 at 315 K, which resulted in the melting and misfolding of H405-411(33) (for more information see Supplementary Material, Supplementary Table 1 and footnotes).

### 2.2 Molecular dynamics simulations

Molecular dynamics calculations that we used in the current work are very similar (except for the SMD part) to the calculations used in our recent work on thermal denaturation of firefly luciferase and are described in greater detail there (33). The structures of the DnaK-Fluc complexes generated by AF3 were solvated 0.15M NaCl was added and the structures energy was minimized for 5,000 steps at 2 fs/step. The complexes investigated were equilibrated at 298 K for a minimum of 200 ns, to relax the AF3 structures. To thermally melt H405-411 in model M3, we set up 15 replicas of the system and ran MD equilibration at 315 K. In one of the replicas H405-411 unfolded within the first 100 ns, after which we lowered the temperature to 298 K and continued our observations of H405-411 and the DnaK lid until both peptides made contact. The isobaric-isothermal NPT ensemble was used in all simulations. For temperature control at 298 K, and 315K, a Langevin thermostat was used with a damping coefficient of 1 ps⁻¹. For pressure control, a Langevin piston barostat was used with a period of 100 fs, decay of 50 fs, and reference pressure of 1.01325 bar (1 atm). NAMD3 (http://www.ks.uiuc.edu/Research/namd/) (34,35) was used with the CHARMM36m force field (36) and TIP3P water model (37). VMD (38) was used to analyze MD trajectories (39) to determine hydrogen bonds between Fluc’s region 403-411 and DnaK region 540-616, and to calculate distances between Fluc’s Ala410 (Model M3) or Thr408 (Model 6) and Met 249 and between Ala410 or Thr408 and DnaK Glu552, as well as to render images of protein structures. Molecular movies were produced based on VMD snapshots using Debut Video Capture Software (NCH Software).

#### 2.2.1 SMD-assisted movement of the alpha helical lid of DnaK

To simulate the movement of the DnaK alpha helical lid that follows the hydrolysis of ATP by NBD during the real chaperone cycle, we applied SMD forces to the Cα atoms of residues 555, 560, and 596, which are located within the helical bundle of the lid. Choosing 3 SMD atoms at different helices, instead of just one atom, applies the SMD potential to the center of mass of that group of atoms, minimizing possible deformations within the lid that could be caused by the external SMD force. To prevent the translation of the system due to external force, the Cα atom of Fluc’s Met249 was constrained in XYZ. The SMD velocity in most simulations involving lid movement was 4×10^-5^ Å/step and in additional simulations testing the dependance of the interactions between the DnaK lid and H405-411, that velocity was decreased 10 times (15 replicas) and 57 times (5 replicas) (Supplementary Figure 6).

#### 2.2.2 Molecular dynamics simulations: sampling procedures

Individual MD simulations (time trajectories) of DnaK lid engaging the Fluc helix differ from each other. These differences are quite significant in the case of the melted Fluc helix, revealing the highly stochastic nature of lid-helix interactions during lid movement. To ensure adequate sampling of this system during the highly dynamic process, we carried out 23 independent replica simulations for the melted helix cases and 28 independent replicas for the intact helix case. In these ensembles, each replica simulation starts from the same initial coordinates (for each system, respectively). However, because we use a Langevin thermostat to control the temperature of the system, atom velocities are randomized at the onset of each replica due to the addition of random Langevin forces at each simulation step, guaranteeing that each replica in these ensembles starts with different velocities and each trajectory is unique.

## 3. RESULTS

### 3.1 AF3-generated DnaK-Fluc complexes

Reconstructing 3D structures of protein complexes when one or more partners exhibit significant conformational dynamics, as is clearly the case with DnaK, remains challenging, both experimentally and computationally (40,41). To visualize plausible complexes of DnaK with Fluc, we used AF3 (32) and followed recently-published approaches aimed at modeling difficult targets that exhibit multiple configurations (41). These strategies include large-scale model sampling (40), generating and comparing models with and without ligands, ions, and including/excluding MSA information (42). Under conditions specified in Materials and Methods and Supplementary Table 1, we generated over 15,000 DnaK-Fluc models. Their detailed analysis is still in process, and the results and our observations, upon completion, will be published in a separate paper. However, we already inspected the highest-ranking models, with a pTM score >= 0.5 (indicative of the global structure being likely correct), which constituted ∼10% of all models. We group them into three distinct classes/clusters. The representatives for each class that received the highest scores are shown in Fig. 1 *ABC* (also Supplementary Table 1).

#### 3.1.1. Cluster 1

Figure 1 *A* shows the highest scoring representative of cluster one (ipTM = 0.72, pTM = 0.74), the highest scoring model globally for all models generated in this work. In this model (Model M1, Supplementary Table 1), DnaK with ATP bound is in the closed conformation, similar to the one reported in (43) (PDB:7kzi), where DnaK, with a short model NR peptide, was captured in an intermediate state before ATP hydrolysis. Fluc is positioned in these models at the bottom of the NBD and makes no contact with the SBD. This group of models seems to provide the tightest interface between Fluc and DnaK, as judged by their ipTM scores, but it is difficult to link such structures to the DnaK chaperone cycle. We speculate that they may represent some Fluc docking configurations that later could develop into structures where Fluc becomes associated with SBD. Although our analysis is not complete yet, it is also possible that these models may be biased by a greater representation of PDB structures in the AF3 internal template set representing the closed conformation of DnaK (either with ADP or ATP), such as 2kho, 7kzi, 7krw, compared to one PDB model with the open conformation (5nro).

#### 3.1.2. Cluster 2

Figure 1 *B* shows the highest-ranking model of this group (model M2, Supplementary Table 1) with DnaK in the ATP-bound open conformation, similar to those reported in (14) (PDB:4B9Q) and in (16) (PDB:4JNE). This specific model was generated with a custom template for DnaK, 4B9Q, and our own template for Fluc, and AF3 was instructed not to use the MSA information (Supplementary Table 1). In M2, Fluc’s structure interfaces with the SBDβ domain in the configuration expected of unfolded proteins interacting with DnaK, however none of Fluc’s structural elements enter the binding cavity. In addition, H405-411, which in this model remains in its unfolded state (thanks to the use of our custom Fluc template, see Material and Methods and Table S1) is away from SBDβ. To further examine this model, we carried out equilibration MD simulations at 298 K to check if Fluc may possibly roll onto the SBDβ into a position that would bring H405-411 closer to the binding site, but the MD results show that the mobility of Fluc and SBDβ in this configuration is limited (data not shown). It is possible that upon ATP hydrolysis, when the lid moves away from the NBD to a position somewhere over the SBDβ, it could come into contact with the misfolded H405-411 and later, during lid opening, could exert some enthalpic pulling forces on it. However, such a scenario was already found to be unlikely (12). Nevertheless, these models warrant some further exploration in future work, but to proceed, it will be necessary to produce many AF3 models of complexes between misfolded Fluc and DnaK in the post ATP-hydrolysis state, with ADP-bound in the NBD.

#### 3.1.3 Cluster 3

In cluster 3, the models present DnaK in a similar conformation as in cluster 2 (4B9Q-like), however, Fluc is placed directly under the DnaK lid, next to the NBD, and not in contact with the SBDβ. The highest-ranking model (model M3, Supplementary Table 1) is shown in Fig. 1 *C*, and Supplementary Figure 2 overlaps this DnaK structure with 4B9Q. The 405-411 helix is positioned around 30 Å away from the αE helix of the DnaK lid. In this model, H405-411 is in the folded state, because this highest-ranking model was generated on the AF3 server, using internal AF3 templates, without our custom Fluc template (model M3, Supplementary Table 1). Similar M3-like configurations were also generated when using our custom Fluc template with H405-411 melted, but those models fell below our pTM cutoff (e.g. model M5, Supplementary Table 1 and Supplementary Fig. 3), so those models were not processed further.

We note that when including our custom template with misfolded Fluc, AF3 frequently “overrode” this template and many models generated with the custom template in the input json file ended up having H405-411 in the folded state. This observation applies equally to M2-like and M3-like models (inspect and compare model M4, and M5, Supplementary Table 1 and Supplementary Figure 3.)

### 3.2. DnaK-Fluc-DnaJ

We wondered whether the unexpected configurations of DnaK-Fluc complexes in M1-like and M3-like clusters are the result of the omission of the crucial DnaK co-chaperone, DnaJ, that is known to mediate the binding of Fluc to DnaK. To test this possibility, we generated 780 DnaK-Fluc-DnaJ models. The results were negative. Like dimers, the trimeric models could be divided into 3 distinct clusters, with the configurations of DnaK and Fluc essentially the same as in the M1-like, M2-like and M3-like clusters without DnaJ (Models M7, M8 and M9, Supplemental Table 1 and Supplemental Figure 4.) suggesting that Fluc location in those models is not influenced by the omission of DnaJ. As in this work we are primarily concerned with events that involve DnaK and Fluc *after* ATP hydrolysis, when DnaJ dissociates from the tertiary complex, we did not include DnaJ when analyzing DnaK’s lid-Fluc interactions (section 3.4).

### 3.3 Limitations of ipTM metrics

The interface predicted template modelling (ipTM) score (44) is used to assess the confidence of AF3-predicted interfaces between polypeptide chains. Scores above 0.8 are thought to indicate high confidence, whereas values below 0.6 suggest low confidence. In our case, model M1 (Fig. 1A) has an ipTM of 0.72, while M2 and M3 score 0.48 and 0.51, respectively (Supplementary Table 1). Examination of M1 shows that its relatively high ipTM value is driven primarily by a small but tight interface between Fluc and the NBD. However, the overall Fluc–DnaK arrangement in M1 does not appear to have immediate mechanistic relevance. In contrast, the global configuration of Fluc and DnaK in M2 aligns more closely with current understanding of DnaK–substrate interactions, yet M2 receives the lowest ipTM score among the three models.

These observations are consistent with recent evaluations of AF3 quality metrics, including ipTM, which highlight their limitations—particularly when multiple heterodimer configurations are possible due to the intrinsic dynamics of one or both partners. In such cases, AF3 models that closely match known PDB structures have nevertheless been assigned unexpectedly low ipTM scores (41). This suggests that current AF3 triaging guidelines may be unreliable for highly dynamic interfaces, which by definition cannot form extensive, stable, and rigid contact surfaces, which would be required to produce high ipTM scores. Because no experimental structures—even partial ones—exist for DnaK–Fluc complexes, this fact underscores the likelihood that these interactions and interfaces must be transient and highly dynamic resulting in an ensemble of complexes rather than a single rigid structure of the two proteins tightly bound together. Thus, any structural models trying to represent this complex biological reality are unlikely to produce very high ipTM scores. The lack of any experimental structural data for DnaK-Fluc complexes that could serve as benchmarks for expected ipTM threshold scores suggests that caution is warranted when evaluating models with ipTM scores lower than suggested by the current guidelines.

For these reasons, we considered not only models with relatively robust ipTM scores such as M1, but also models like M2 and M3 whose lower scores likely reflect the presence of rather weakly bound interfaces that should be expected for such a dynamical system. Despite their lower ipTM values, these models may be biologically relevant, and they certainly provide extremely valuable starting structures for developing mechanistic hypotheses that may be experimentally verified.

More broadly, we view the strength of the AF3 platform for systems with dynamic interfaces—such as DnaK and Fluc—not as a substitute for missing experimental data but as a generator of new conceptual possibilities that would not emerge otherwise. In our case, AF3 uniquely produced three classes of physically plausible DnaK–Fluc models (i.e., models without steric clashes), including configurations that are counterintuitive from the perspective of existing structural intuition. Importantly, that intuition has been shaped almost entirely by structures of DnaK bound to very small peptides rather than to real, large substrates such as Fluc. AF3, by contrast, is not constrained by the limitations of peptide-based model systems. The conceptual avenues opened by these models can, in turn, motivate new experimental strategies aimed at testing and refining the hypotheses they suggest.

### 3.4 Interactions of DnaK lid with H405-411 in the M3-like complex

Our expectations that the DnaK lid may interact with Fluc’s helix 405-411 in M3-like configurations, are rooted in preliminary computational experiments with the very first model of the DnaK-Fluc complex that we generated with AF3, shortly after its release in May 2024. In this model (now coined M6, Supplementary Table 1, Supplementary Figure 5), Fluc’s H405-411 was positioned directly under the SBDα kink and molecular dynamics simulations captured very strong hydrogen bonding between DnaK lid residues such as Gln549 and Glu552 and Fluc’s residues Asn409 and Thr408, respectively. However, in the current highest-ranking model M3, H405-411 is further away from the lid, as compared to M6, so it is not obvious, whether similar interactions between the DnaK lid and the helix could be captured for M3.

To test if these interactions between DnaK and Fluc would occur in M3, we used the predicted atomic coordinates of M3 to build an all-atom model of the solvated complex for molecular dynamics simulations (Materials and Methods), allowing us to observe the complex’s dynamics through a “computational microscope” (45,46).

#### 3.4.1 Interactions of the DnaK lid with the misfolded H405-411

We started by equilibrating the DnaK-Fluc complex at 25 °C for 200 ns. Then, to melt H405-411 we raised the MD temperature to 42 °C. As our recent MD study of *isolated* Fluc indicated that it takes on average 2.5 microseconds to melt H405-411 at 315 K *in silico* (31), we set up 15 replicas of the DnaK-Fluc complex at 42 °C to increase our chances to capture helix melting earlier. Indeed, in one of the replicas, the helix melted after only 100 ns and we arrested this misfolded structure by lowering the temperature back to 25 °C. We continued the equilibration at 25 °C to see if the DnaK lid would approach the melted H405-411. This “search” process, which initially is completely random, is illustrated as a molecular movie in Supplementary Fig.2.360 that accompanies Fig. 2 in the main text. It is quite striking to observe that after a long random walk, when the lid finally approaches the helix, the interactions between the two become so strong that the helix “jumps” to contact the lid (Fig2.360, t= 25 s into the movie). These results directly show that in the initial complex the melted helix of Fluc does not need to be in the immediate vicinity of the DnaK lid to be eventually recognized by the lid. This observation in turn supports the idea expressed in point 3.3 above that an ensemble of DnaK-Fluc complexes rather than a single structure is expected at early stages of DnaK-Fluc interactions and that the modest tightness of the interface between Fluc and DnaK is actually advantageous for the search and recognition process.

**Figure 2.**
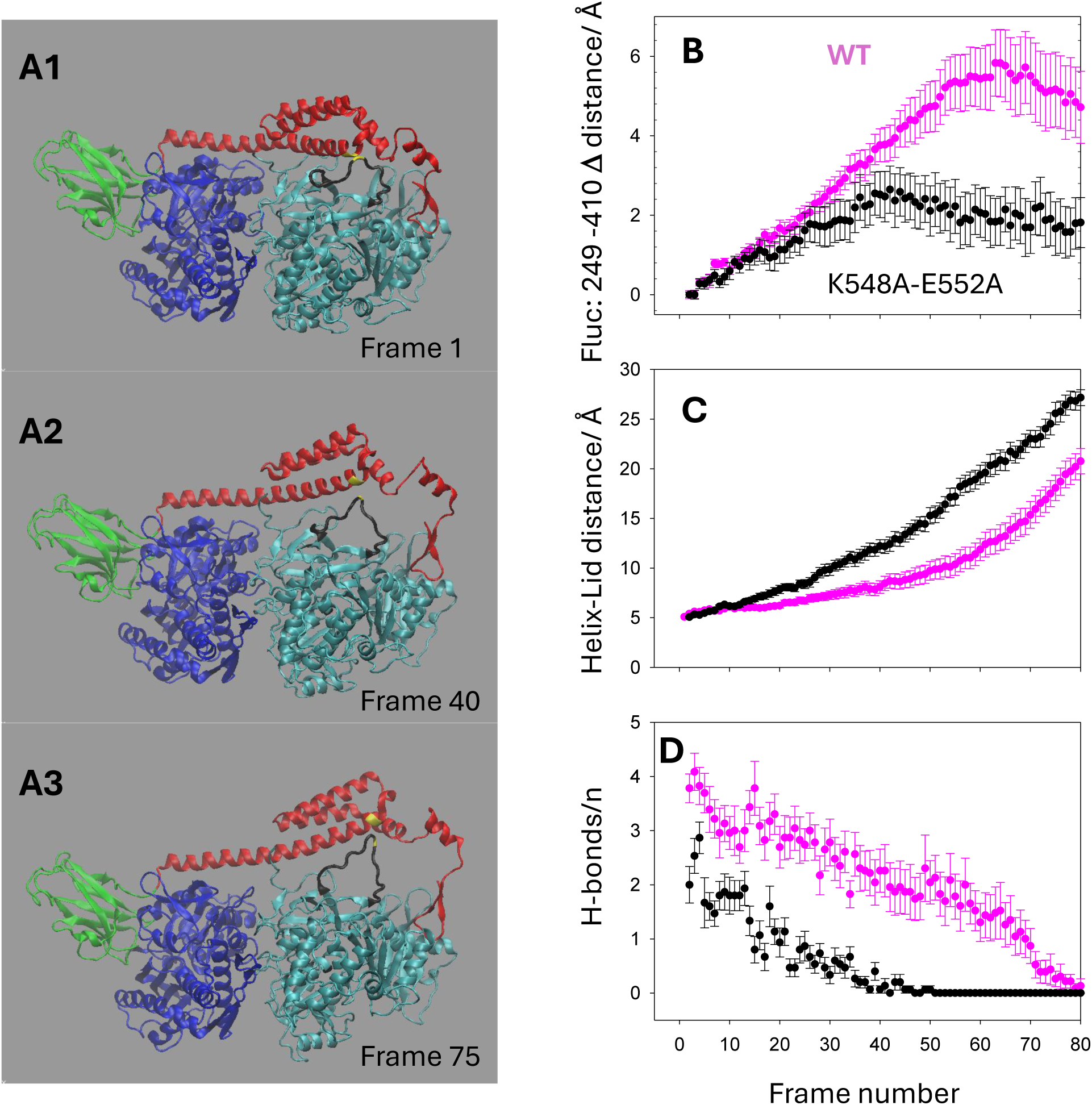
Steered molecular dynamics simulations capture the interaction between the DnaK lid and Fluc’s H405-411 residues that pull the helix out of Fluc (black ribbon structure represents the whole melted 402-417 region that encompasses H405-411). Three MD snapshots are shown at given MD frames in (A1-3). See also the accompanying 360 movie that shows the MD trajectory revealing the initial “search” process where the lid finds H405-411 followed by a SMD trajectory where the lid is pulling the helix (B) shows the increment in the separation of the helix from the Fluc’s center as measured by the changes in the Met249-A410 distance. (C) shows the distance between Ala410 of the helix and Glu552 of the lid. (D) shows the number of hydrogen bonds between Fluc’s region encompassing residues 403-413 and Dnak lid’s residues 540 to 616. H-bonds were determined for each MD frame in VMD. The purple points in B, C and D represent the data for the WT DnaK, while the black points represent the data for the double mutant of the DnaK lid.

After the lid engaged H405-411, we forced the lid to move gradually away from Fluc towards the SBDβ, to simulate the known lid behavior that follows ATP hydrolysis by NBD when the lid moves to the closed conformation (14,16). Briefly, SMD forces were applied to the DnaK lid through Cα atoms of residues 555, 560 and 596 while the Cα atom of Fluc’s Met249 was constrained in space to avoid DnaK-Fluc translocation (Materials and Methods).

In Fig. 2A, we show the results of one of 23 replica simulations involving DnaK lid movement. MD snapshots were captured at the onset of the movement of the lid (A1), in the middle (A2) and shortly before the contact between the lid and the helix broke (A3). It is clear, based even on a simple visual inspection of this MD trajectory, that H405-411 follows the movement of the lid and is effectively pulled out of the Fluc structure. The MD-based animation of this experiment is included in the movie in Fig. 2.360 (last 10 seconds), and more examples are available as molecular movies in the Supplementary Material. It must be emphasized that in these simulations, the external SMD force is applied only to the alpha helical lid of DnaK to guide it toward the SBDβ and not to Fluc. The only external forces that Fluc and its H405-411 experience in this system come from the interactions with DnaK (and with the solvent). To quantify the displacement of H405-411relative to the Fluc core, when pulled out by the DnaK lid, we measured the distance of Ala410 from Fluc Met249, which is buried close to the center of Fluc, its position does not fluctuate widely. We chose Ala410 as our focal residue, because this is the helix residue that “jumps” to contact Glu 552 in the DnaK lid, and because it is close to the middle of the whole melted region 402-417 that encompasses H405-411. The increment of this distance, Δ, *averaged* over 23 independent replica MD trajectories, is shown in Figure 2B, as a pink trace. To quantify the position of the melted helix, relative to the DnaK lid during the lid movement, we measured the distance between Fluc Ala 410 and DnaK Glu552, which is located right at the kink of the lid and is very close to Ala410 at the onset of lid movement. This distance is shown in Fig. 2C as a pink trace, and this distance remains small (5-7 Å) and quite constant for the first 40 MD frames, while the melted helix and its neighboring residues closely follow the lid. To investigate the origin of the interactions between the melted region and the DnaK lid that allow this pulling process to occur, we examined hydrogen bonds between the two structures (see Materials and Methods for details) during the MD simulations. The average number of hydrogen bonds between the lid and the helix region are plotted in Fig. 2C, as pink symbols. These bonds, analyzed in VMD (38), include side chain - side chain and side chain - main chain interactions indicative of intimate interactions between the lid and the helix. The bonds’ occupancy is tabulated in Supplementary Table 2. Asp413-Lys548 and Ala410-Glu552 are among the most persistent bonds during lid movement. The overall timing of the movement of the DnaK lid from its open to closed conformation is likely related to the timing of the chaperone cycle and the rate of ATP hydrolysis. However, it is presently unknown what the actual velocity of the lid at the onset of that transition may be. Our SMD pulling velocity of 4x×0^-5^ Å/step (2 m/s) applied to the lid in the simulations shown in Fig. 2 is in the range of SMD velocities typically used when modeling intermolecular hydrogen bonds under mechanical forces (47). However, the results of such non-equilibrium transitions may generally depend on the speed at which forces are applied to these hydrogen bonds (47). To test the sensitivity of our results to varying lid velocity, we repeated these simulations at a 10 times slower speed (15 replicas) and at a 57 times slower speed (5 replicas). The results for the three speeds tested are shown together in Supplementary Figure 6.

Three molecular movies (one for each velocity) are shown in the Supplementary Material. We did not observe any significant changes in the lid-helix behavior in the range of velocities tested.

#### 3.4.2 DnaK lid mutants

To verify that the hydrogen bonds between the lid and H405-411 are key to the observed helix pullout, we mutated *in silico*, the key lid residues. First, we replaced Glu552 with alanine, but this single point mutation had only a minor effect, and the helix was still able to follow the lid during its movement away from Fluc (data not shown). We then created a double mutant E552A; K548A and this mutation had a strong detrimental effect on the ability of the lid to pull the helix and its flanking residues (Fig. 2 *BCD*, black symbols), although it did not abolish the helix movement completely. We investigated the hydrogen bonding pattern between the mutant and the helix in those simulations and found that 1-3 bonds during the early stage of the simulation are still maintained (Supplementary Table 3), which is enough for a small helix displacement.

#### 3.4.3 Interactions between the DnaK lid and the intact H405-411

To examine these interactions, we used our first AF3 model of the DnaK-Fluc complex, M6, which is in an M3-like configuration. We chose M6 (Supplementary Table 1, Supplementary Figure 5), because in this model the Fluc orientation relative to SBD*α* is such that the *intact* H405-411 is already under the DnaK lid. We also melted H405-411 in M6 (Supplementary Figure 7) so we could compare the behavior of the intact helix and the melted helix using the same model, M6. We performed SMD simulations, on M6 in both cases similar to those described above, with the DnaK lid guided toward SBDβ by SMD forces. The snapshots of the complex from these simulations are shown in Supplementary Figure 8 *AB*, and MD trajectories are shown as molecular movies in the Supporting Material. The results of 28 independent replica simulations with the intact helix and of 23 replicas for the melted helix are averaged and summarized in Supplementary Figure 9 *ABC*. From Supplementary Figure 8 *B*, it is evident that when the helix is intact, it does not follow the DnaK lid during its movement toward SBDβ and it stays close to its original position within the Fluc structure. This indicates that any pulling forces exerted by the lid on the helix cannot overcome the restoring forces that keep the helix folded and in place. As shown in Supplementary Figure 9, both metrics used to evaluate the position of the intact helix 405-411 relative to the DnaK lid and the Fluc core are significantly different as compared to the data for the melted helix. Also, the behavior of the hydrogen bonds between the lid and the helix during lid movement is very different in both cases. The bonds to the intact helix are broken much earlier in the process as compared to the persistent bonds to the melted helix that facilitate its pullout.

### 3.5 Helix refolding

It is tempting to speculate that pulling out the misfolded helix from the Fluc surface by the DnaK lid, helps the helix overcome the energy barriers that it faces on its path to the native state, accelerating its refolding. To examine this possibility, we carried out 10 relaxation simulations to follow the helix behavior, after it breaks off from the DnaK lid.

In one of these simulations, after approximately 3 μs, the 402-417 region, encompassing the helix, formed a one turn folding intermediate, while in the other 9 replicas this region remained misfolded. Because of the computational cost, we abandoned those not-so-promising simulations and continued the simulation that harbored the folding intermediate. The results are shown in Fig.3 and its accompanying animation (Fig 3.360). For the next ∼7 μs the intermediate occasionally reversed to the misfolded state but did not cross the barrier to the native state. However, at around 10.6 μs, the second helical turn was formed suddenly, and after a short instability (inspect the animation) it formed a stable 2 turn native helix that persisted for the next 533 ns, until the end of our simulation. This result is promising, as in our previous work with the melted 405-411 helix we did not observe any *spontaneous* refolding at 298 K, for the total simulation time of 18 μs (33). However, more simulations will be required to examine, whether the average refolding time of H405-411, following the DnaK lid-driven pullout, is significantly shorter (in the statistical sense), as compared to any spontaneous refolding.

**Figure 3.**
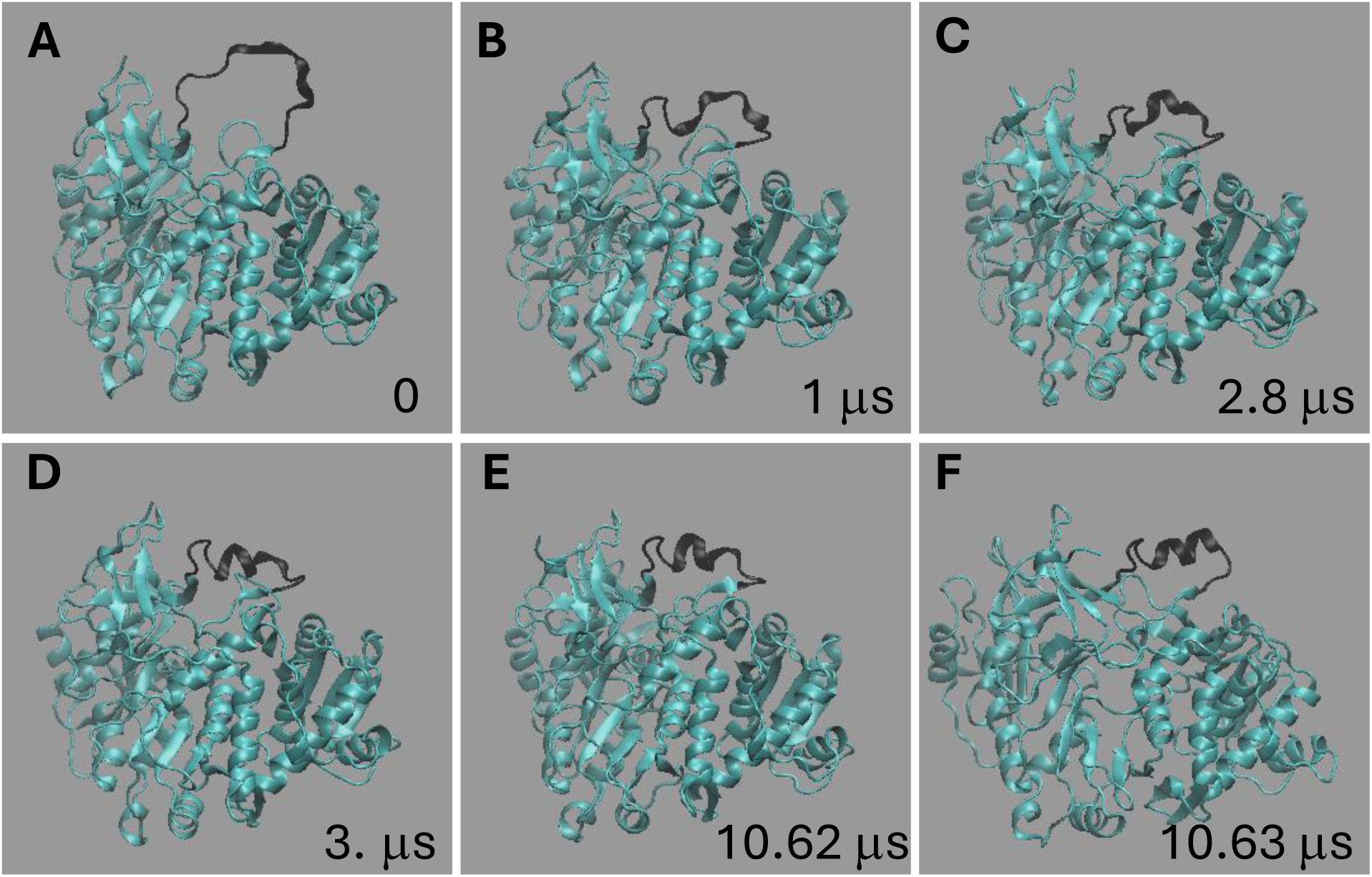
Snapshots of Fluc MD relaxation simulation after H405411 detached from the DnaK lid following its pullout from the Fluc surface capture H405-411 complete refolding to the native state. See also the accompanying 360 movie.

## 4. DISCUSSION

Using AF3, we generated many models of the complex between DnaK chaperone and Fluc in its native and misfolded states that clustered into three distinct groups. We examined the models with the highest confidence scores and evaluated their relevance for the chaperone mechanism. Model M1 received the best metrics but its biological relevance is at present unclear. Model M2 places Fluc close to the SBDβ part of the substrate binding domain, as expected based on the studies with short peptides or substrates that have large unstructured regions. However, it is unclear how DnaK could refold Fluc’s H405-411 in this configuration. It might be possible that SBDβ could engage Fluc if it had large misfolded and extended regions accessible to the SBDβ cradle; however, there is no experimental evidence that such regions do exist in Fluc at 42 °C. The other possibility is that after ATP hydrolysis, the DnaK lid could contact Fluc when the lid returns to the closed conformation, possibly then interacting with H405-411, but structures that capture such configurations are not yet available. On the other hand, in model M3, Fluc is placed directly under the SBDα part of the substrate binding domain but does not contact SBDβ. This arrangement provides a quite compelling new scenario for how DnaK might interact with Fluc during its chaperone action as its alpha helical lid is already in intimate contact with Fluc, and because of inherent lid mobility the lid may get close to misfolded elements present in Fluc *before* ATP hydrolysis.

To examine the interaction between DnaK and Fluc in M3-like configurations in detail, we focused “our computational microscope” on the interface between the DnaK lid and the Fluc region 402-417, encompassing H405-411. We melted the 405-411 helix at 42 °C and equilibrated the system at 25 °C while observing the behavior of the melted helix and the DnaK lid. At first the lid moves randomly, but when by chance approaches the melted helix, the interactions between the two become evident as the helix “jumps out” to contact the lid. Then we simulated the movement of the DnaK lid toward the SBDβ and observed the behavior of the melted 402-417 region of Fluc. We observed that the melted helix within the 402-417 melted region, follow the DnaK lid closely.

These residues are effectively pulled out of the Fluc structure and stretched. When we mutated (*in silico*) the key lid residues K548 and E552, which form persistent hydrogen bonds with the helix region, the displacement of the helix was significantly diminished. When the DnaK lid interacted with the intact H405-411 and was moved towards SBDβ the intact helix did not follow the DnaK lid and remained within the Fluc fold. When the helix is intact, it is quite rigid and not very deformable; thus, the hydrogen bonds with the lid are easily broken upon lid departure, and there is not enough force to deform and pull out the helix. Importantly, we did not observe the folded helix to “jump out” to contact the lid, which may suggest that the lid does not engage folded elements that do not present themselves for interactions with the lid. It is the melted helix’s flexibility that allows it to form persistent hydrogen bonds with the lid and thus the pulling force to persist (up to a point when the elastic restoring forces exceed the strength of these hydrogen bonds). As a result, the melted helix can “jump” to contact with the lid and can be pulled out of its deep local minimum energy state (the misfolded state), allowing it to refold upon detachment from the lid and the relaxation due to the restoring forces.

The most significant advantage of this new model is that it provides a direct connection between ATP hydrolysis by NBD and a mechanical action of the chaperone on its substrate. We show that DnaK does not need to spend any ATP-derived energy to identify misfolded elements in its substrates. The misfolded elements would identify themselves, by “jumping out” towards the DnaK lid when its nearby. The energy produced by ATP hydrolysis would be used to pull those elements further away from the substrate, which would convert a part of this energy into elastic energy stored in the stretched segment. This extra energy could potentially help to cross the energy barrier on the path to the native state.

It is impossible to claim with certainty the placement of Fluc as in M3 represents biological reality for this complex. However, it is possible that DnaK may interact differently with small model peptides such as NR or proteins that have intrinsically unstructured regions such as σ^32^ (12) or proPhoA (13) and differently with large, well-structured substrates such as Fluc. We hypothesize that such a distinction may be related to the various DnaK functions present when interacting with these structurally diverse substrates. The primary function of DnaK when interacting with the σ^32^ regulatory protein is to capture and sequester it, while with proPhoA, DnaK binds to it and prevents its premature folding. In both cases DnaK does not need to mechanically remodel these proteins. However, when interacting with denatured Fluc, DnaK needs to restore its correct fold. This may occur by a mechanical action involving the extraction of the misfolded elements out from their low energy misfolded states, while consuming the ATP energy. Thus, our observations do not question the large body of work on the DnaK mechanism when the chaperone interacts with unstructured proteins. Instead, we suggest some new possibilities for the DnaK assisted protein refolding mechanism.

Although we do not provide any experimental evidence for this new model, our observations are consistent with DnaK lid truncation studies, which showed that removing the C terminal part of the lid up to around position 603 (16,48) does not affect DnaK ability to refold denatured Fluc, however, removing the lid fragment up to position 538, abolishes its chaperone activity (49). We provide a plausible new interpretation of these results. When the lid is truncated at position 538, all critical residues that are expected to form hydrogen bonds with Fluc are eliminated, so the lid cannot pull misfolded structures out. The large truncation certainly affects also the binding and retention of intrinsically unfolded substrates in the SBDβ, so there is no contradiction between the two models when applied to appropriate substrates. In addition our observations are also consistent with studies that determined that immobilization of the lid by covalently connecting it to the SBDβ (via di-sulfide bonds) in the closed conformation, abolishes DnaK’s ability to refold Fluc (50). Moreover, the ability of DnaK to resolve small aggregates, either alone or together with ClpB (7) seems to be also consistent with our model, which would allow DnaK to engage larger structures by throwing the SBDα lid on them and “fishing out” misfolded elements. Finally, we propose an experiment that could verify our conjectures. It would be important to determine where is thermally denatured Fluc located in relation to SBDβ while in a complex with DnaK. Such a determination should not, in principle, need to involve high resolution structural studies, which have proved very difficult for that complex, so far. Well-designed SM FRET measurements could possibly provide the answer.

## 5. CONCLUSIONS

Using computational approaches, we investigated the interactions between DnaK chaperone and firefly luciferase in its native and denatured states. We propose that Fluc is associated with the DnaK lid (SBDα) and the DnaK nucleotide binding domain (NBD), rather than with its substrate binding domain, SBDβ. The DnaK lid forms labile hydrogen bonds with Fluc’s melted helices and pulls them out from the Fluc structure during ATP-hydrolysis-powered movement of the lid away from the substrate. Melted helices, which are forcefully pulled out and stretched, can refold upon relaxation, while intact helices are not disturbed. These observations, although speculative at present, may lay the foundation for a new model of the DnaK-assisted protein refolding mechanism.

## Supporting information

Supplementary tables and seven supplementary movies

## Acknowledgments

PEM thanks Monika Marszalek for her help with data processing and figure preparation, and Tom Pan for cloning and setting up the GitHub AF3 version 3 on the Duke University Computer Cluster. PEM acknowledges Duke Research Computing for access to the Duke Computer Cluster. This work was supported in part by the National Science Foundation grant MCB 2118357.

## Author contributions

PEM: conceptualization, executing and analyzing molecular dynamics simulations, writing, and funding acquisition. OKM: submitting AF3 jobs and analyzing AF3 outputs, evaluating AF3 models and confidence metrics, manuscript editing and revisions.

