## Supplementary tables and seven supplementary movies for "DnaK refolds denatured proteins by actively pulling out their misfolded structural elements": DnaK-2026-Supplementary-Material-rev1.pdf

**Supplementary Material description**: Three supplementary tables, 9 supplementary figures and 7 supplementary movies.

### Supplementary Material

### Running AF3 jobs on AF3 server and locally (Duke Univ. computer cluster, DCC)

**AF3 server.** We used the official DeepMind AF3 WEB-based interface and submitted up to 30 jobs daily, using random seeds and specifications as indicated in the table below. AF3 version 1 (V1) was used.

**DCC.** When submitting batch jobs on the Duke University computing cluster, we typically specified up to 50 unique seeds/per job and on the days when the cluster was less busy, we could run up to 10 jobs per night, generating up to 2500 models a day. AF3 output for each job includes an Excel table ranking all models within a job (up to 250 models/job in our case), making the initial screening of models relatively simple. For in- depth structural analysis we chose models with pTM scores >=0.5 (with some exceptions, e.g. models M8 and M9 for DnaK-Fluc-DnaJ complexes). AF3 version 3 (V3) was used.

**Supplementary Table 1. AF3-generated Fluc-DnaK models used in this work**

| # | Model "code" | Clu ster | ipTM | pTM | Un-paired MSA | Paired MSA | ATP | Na Cl | AF3 templates | Custom templates | AF3 V1 or V3 |
|---|---|---|---|---|---|---|---|---|---|---|---|
| M 1 | 898M4 | 1 | 0.72 | 0.74 | + | + | + | + | - | - | V3 |
| M 2 | 10010M1 | 2 | 0.48 | 0.52 | - | - | + | + | - | 4B9Q* C2-73-315K** | V3 |
| M 3 | 748M0 | 3 | 0.51 | 0.56 | + | + | + | + | + | - | V1 |
| M 4 | 10007M4 | 2 | 0.47 | 0.52 | - | - | + | + | - | 4B9Q C2-73-315K | V3 |
| M 5 | 10003M2 | 3 | 0.45 | 0.49 | - | - | + | + | - | 4B9Q C2-73-315K | V3 |
| M 6 | May2024 | 3 | 0.24 | 0.51 | + | + | - | - | - | - | V1 |
| M 7 | 86M3 | 1 | 0.57 | 0.63 | + | + | + | + | + | - | V3 |
| M 8 | 98M4 | 2 | 0.42 | 0..47 | + | + | + | + | + | - | V3 |
| M 9 | 199M4 | 3 | 0.41 | 0.44 | + | + | + | + | - | - | V3 |

* PDB: 4B9Q is the critical DnaK structure in an ATP-bound open conformation, which was used as our custom template, as it was not reported among the internal AF3 templates in the AF3 output data json files that we inspected.

** C2-73-315K is our Fluc structure with the H405-411 helix unfolded via a MD simulation at 315 K, which belongs to the data set that we previously reported. We used this structure along with 4B9Q in some AF3 jobs (AF3 version 3).

“Model code” refers to our internal naming methodology for AF3 models produced in this work, which uses the unique AF3 seed number followed by the number of the specific model (0....4) as each seed produces 5 different models. This simple method allows us to quickly identify and retrieve the models of interest among the vast number of all outputs.

**Supplementary Table 2**. Hydrogen bonds between WT-DnaK Lid and Fluc (403-413) during the SMD-controlled lid movement

| donor | acceptor | <occupancy> / % |
|---|---|---|
| LYS548-Side | ASP413-Side | 27.3 |
| ALA410-Main | GLU552-Side | 23.0 |
| ASN409-Side | GLN603-Side | 12.4 |
| LYS548-Side | LEU411-Main | 11.0 |
| THR408-Main | HSD606-Side | 8.7 |
| GLN549-Side | ASN409-Side | 8.3 |
| ASN409-Side | GLU552-Side | 7.0 |
| HSD544-Side | ASP413-Side | 4.1 |
| ASN409-Side | GLN549-Side | 3.8 |
| ASN409-Side | GLN549-Side | 3.5 |

**Supplementary Table 3**. Hydrogen bonds between the double DnaK Lid mutant (Lys548Ala and Glu548Ala) and Fluc (403-413) region during the SMD-controlled lid movement

| donor | acceptor | <occupancy> / % |
|---|---|---|
| GLN549-Side | ASN409-Side | 8.5 |
| THR408-Main | HSD606-Side | 8.0 |
| ASN409-Side | GLN603-Side | 3.7 |
| ASN409-Side | GLN549-Side | 3.1 |
| GLN603-Side | ASN409-Side | 0.8 |
| ASN409-Side | HSD606-Side | 0.5 |
| HSD544-Side | ASP413-Side | 0.3 |
| LEU411-Main | GLU551-Side | 0.3 |
| ALA407-Main | HSD606-Side | 0.3 |

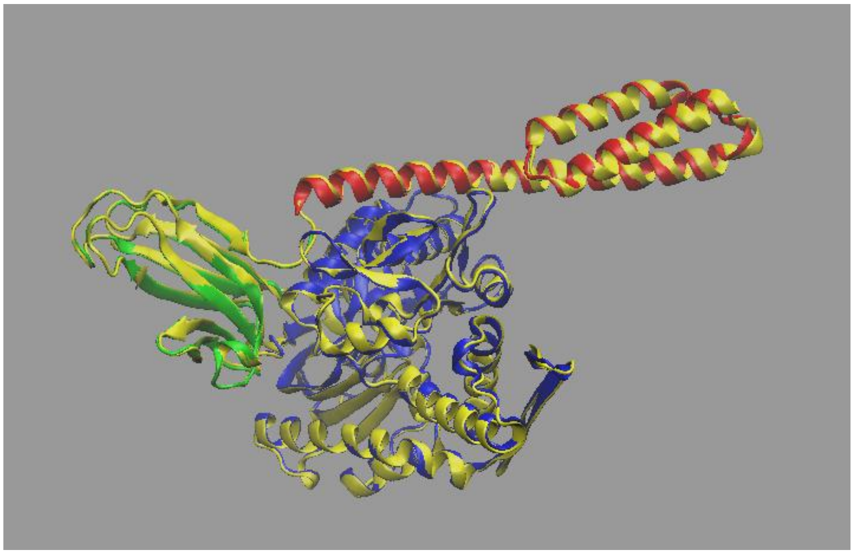

**Supplementary Figure 1.** *Comparing the structure of DnaK in AF3 model M2 with PDB:4B9Q*. The DnaK backbone from 4B9Q is depicted in yellow. The DnaK domains from M2 are coded in the usual colors. The domains were individually aligned with 4B9Q over respective sequences in VMD.

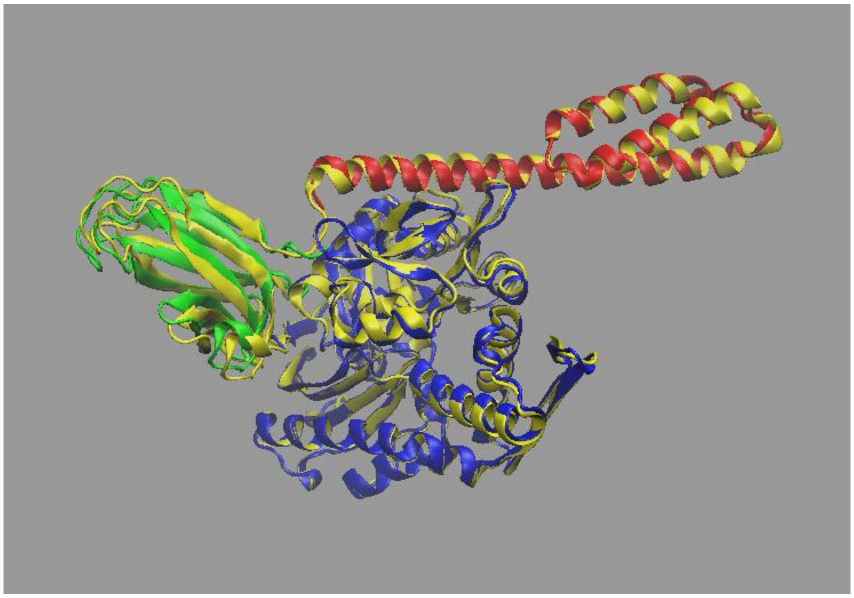

**Supplementary Figure 2*A*.** *Comparing the structure of DnaK in AF3 model M3 with PDB:4B9Q*. The DnaK backbone from 4B9Q is depicted in yellow. The DnaK domains from M2 are coded in the usual colors. The domains were individually aligned with 4B9Q over respective sequences in VMD. The overall RSMD of 1.07 Å was calculated in VMD as the weighted average, where the number of residues in each domain served as weights.

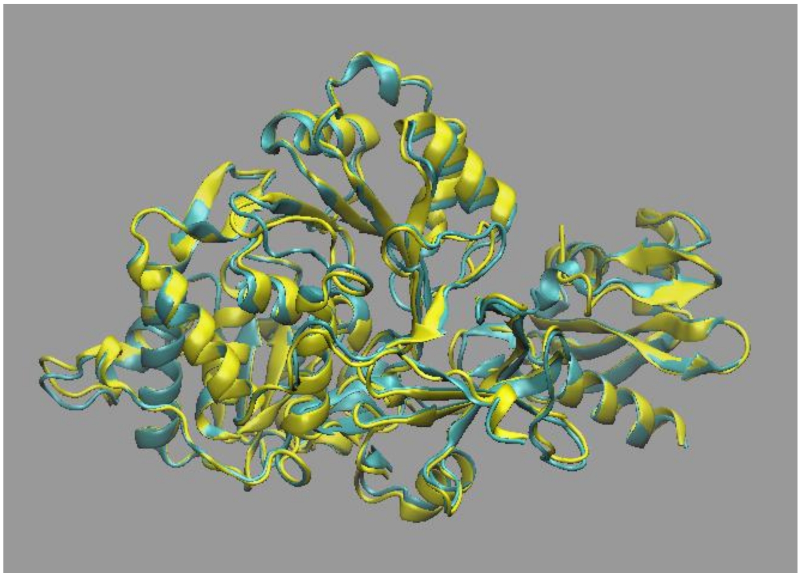

**Supplementary Figure 2*B*.** *Comparing the structure of Fluc in AF3 model M3 with PDB: 1BA3*. Fluc from M3 is shown in cyan and from 1BA3 in yellow. Backbone RMSD = 1.08 Å.

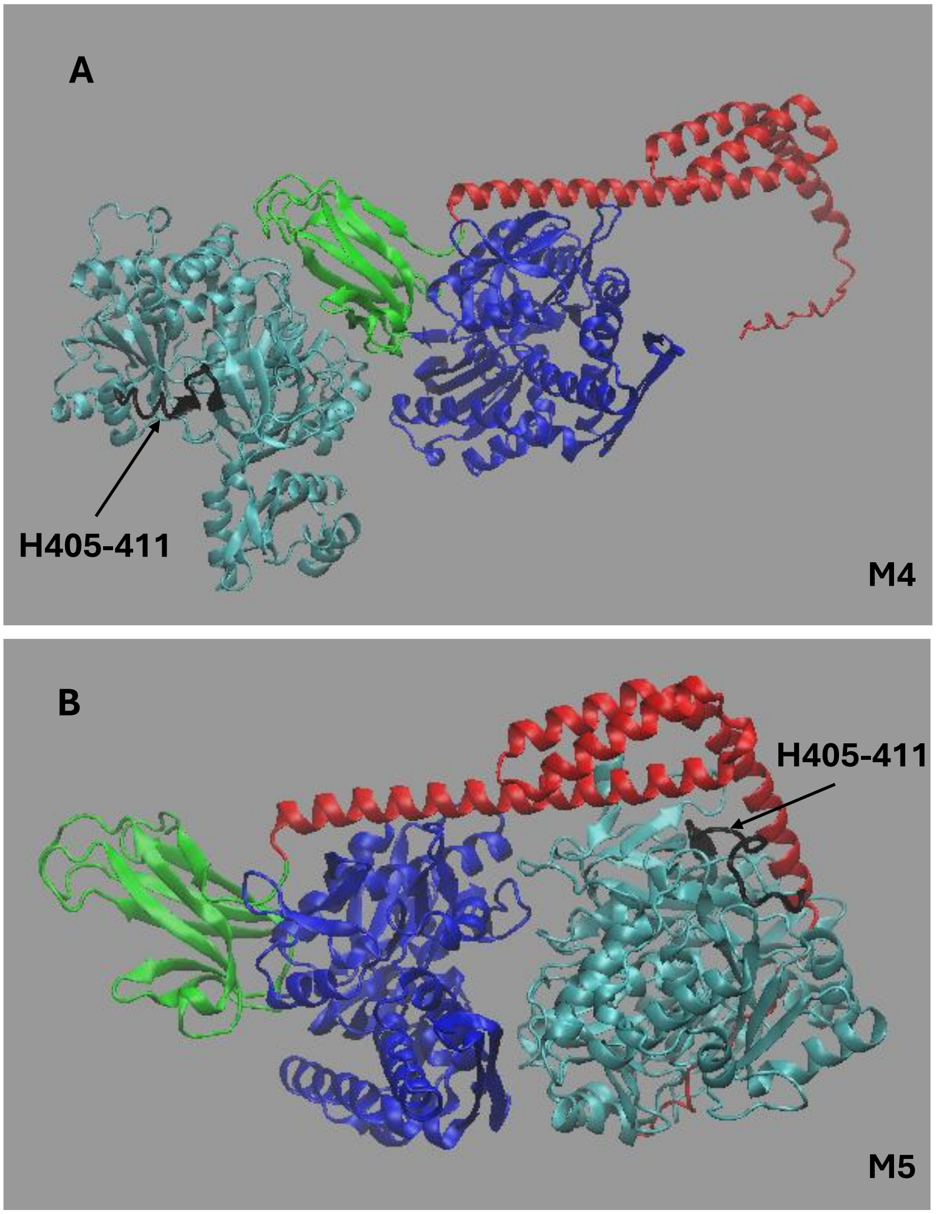

**Supplementary Figure 3***. Examples of AF3 models of DnaK-Fluc complexes produced when using our custom template for Fluc, with H405-411 in the thermally unfolded state*. Under the same modeling conditions (Supplementary Table 1), in M4 (***A***), H404 was produced in the native ***folded*** state (unlike in M2, main text, where it was unfolded), while in M5 ***(B)***, H404-411 was produced in the **unfolded** state.

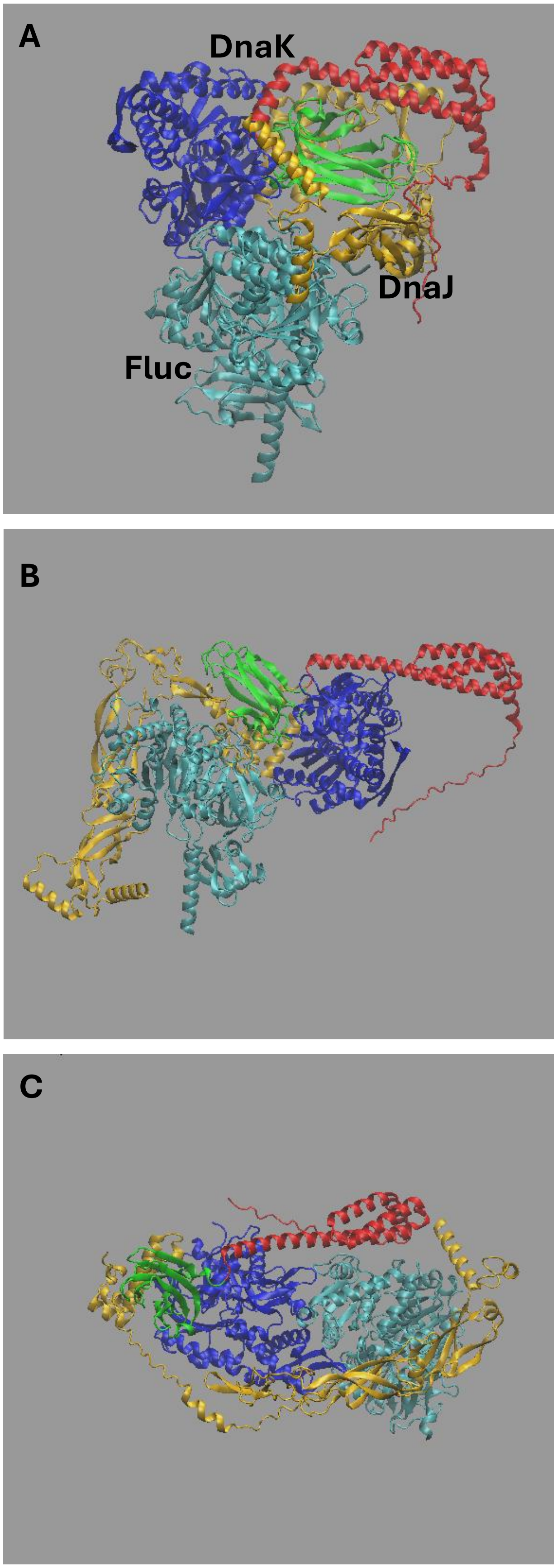
A
DnaK
DnaJ
Fluc
B
C

**Supplementary Figure 4.** *Structures of trimeric complexes of DnaK-Fluc-DnaJ.* DnaJ is shown in gold color**.** Like dimeric DnaK-Fluc complexes, AF3 models group into 3 clusters, with DnaK-Fluc configurations shown in *ABC*, consistent with those in the dimeric models M1, M2 and M3, respectively (Fig. 1, Main text, Supplementary Table 1).

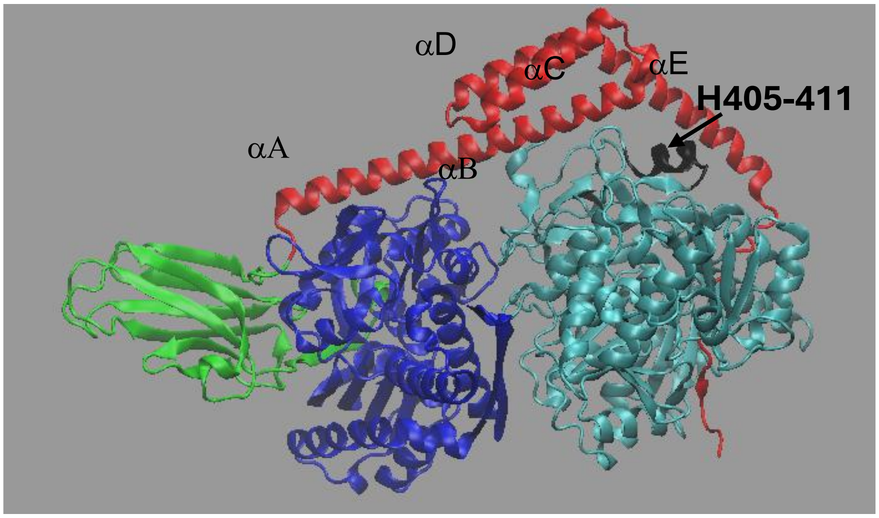

**Supplementary Figure 5.** *Our first AF3 model of DnaK-Fluc.* Generated in May 2024. Now this model is named M6 (Supplementary Table 1), because of its lower ipTM score as compared to the new model M3.

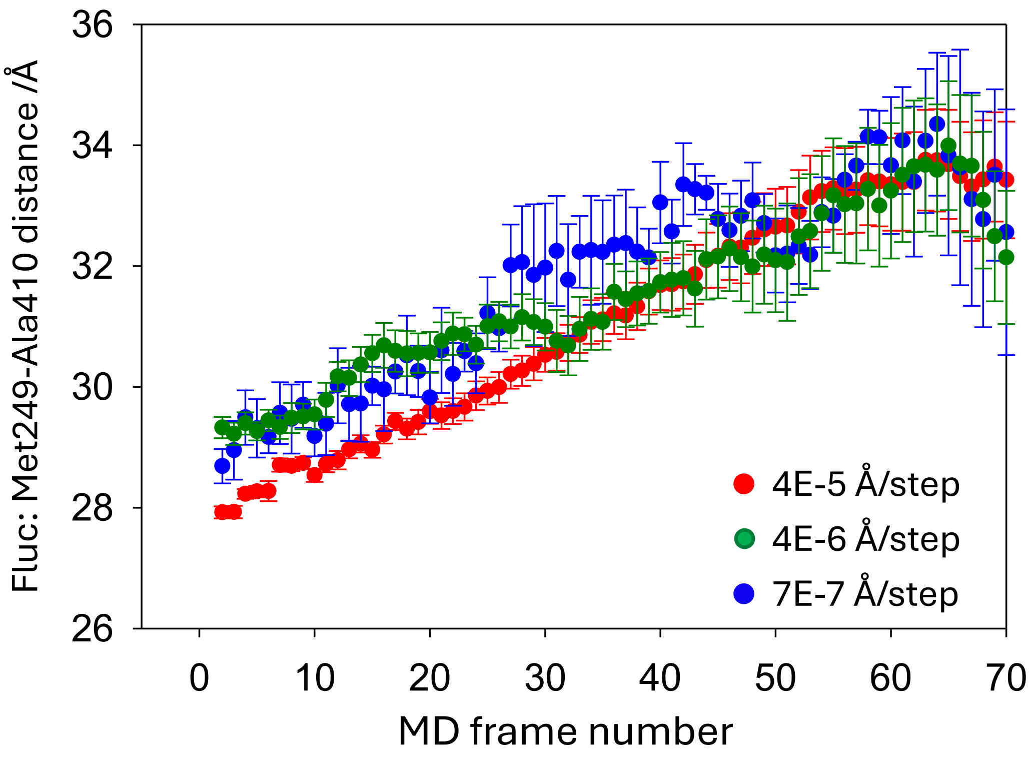

**Supplementary Figure 6.** *DnaK pulls out H405-411 at various lid speeds (Model M3).* The trace shown here in red is the same as the pink trace in the main Fig. 2*B*, except that here the absolute Met249-Ala410 distance is shown and in Fig. 2 the increment is shown.

ns).

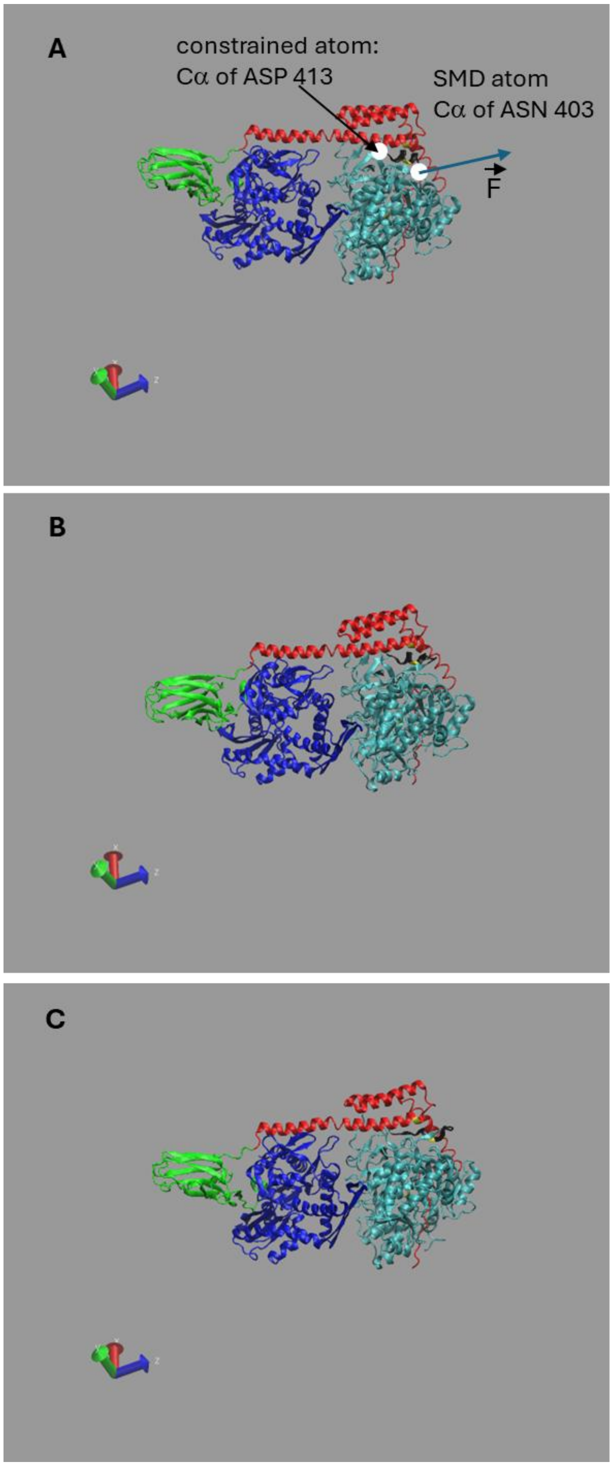

**Supplementary Figure 7. Model M6**. *SMD-stretching of the 405-411 helix within the Fluc segment 403-413, in the DnaK-Fluc complex simulated in all-atom MD calculations at 42 °C (315 K).* **A**, starting frame (t=0); **B**, frame 97 (t = 1.94 ns); **C**, frame 155 (3.1 ns).

The NAMD restart files generated at the 350 ns step of the DnaK-Fluc **model M6** equilibration simulation at 315 K were used to execute a short 3 ns simulation of the whole system aimed at forcing the 405-411 helix to melt. The C$\alpha$ atom of Fluc residue 413 was constrained in XYZ while the C$\alpha$ atom of residue 403 was subjected to a stretching force using the SMD protocol within NAMD. Briefly, in this approach, an external force is applied to the selected SMD atom in the form of a virtual harmonic spring, whose one end is attached to the SMD atom and the other end is moved at a certain, constant velocity in a given direction. In our case, the stretching velocity was $10^{-5}$ Å/step and the SMD direction was (0 0 1). As a result of the applied force, the 405-411 helix melted, and it did not relax back after removing the SMD force and cooling the whole system at 298 K for 115 ns. Of note is that SMD stretching at 298K caused only a reversible extension of the helix, which was not useful. The structure of the complex, with the melted helix generated this way is shown in **Supplementary Figure 8 *A1*** and was used as a starting point for all 23 simulations aimed at following the interaction between the DnaK lid and the melted helix in Model 6, which are averaged in Supplementary Figure 9.

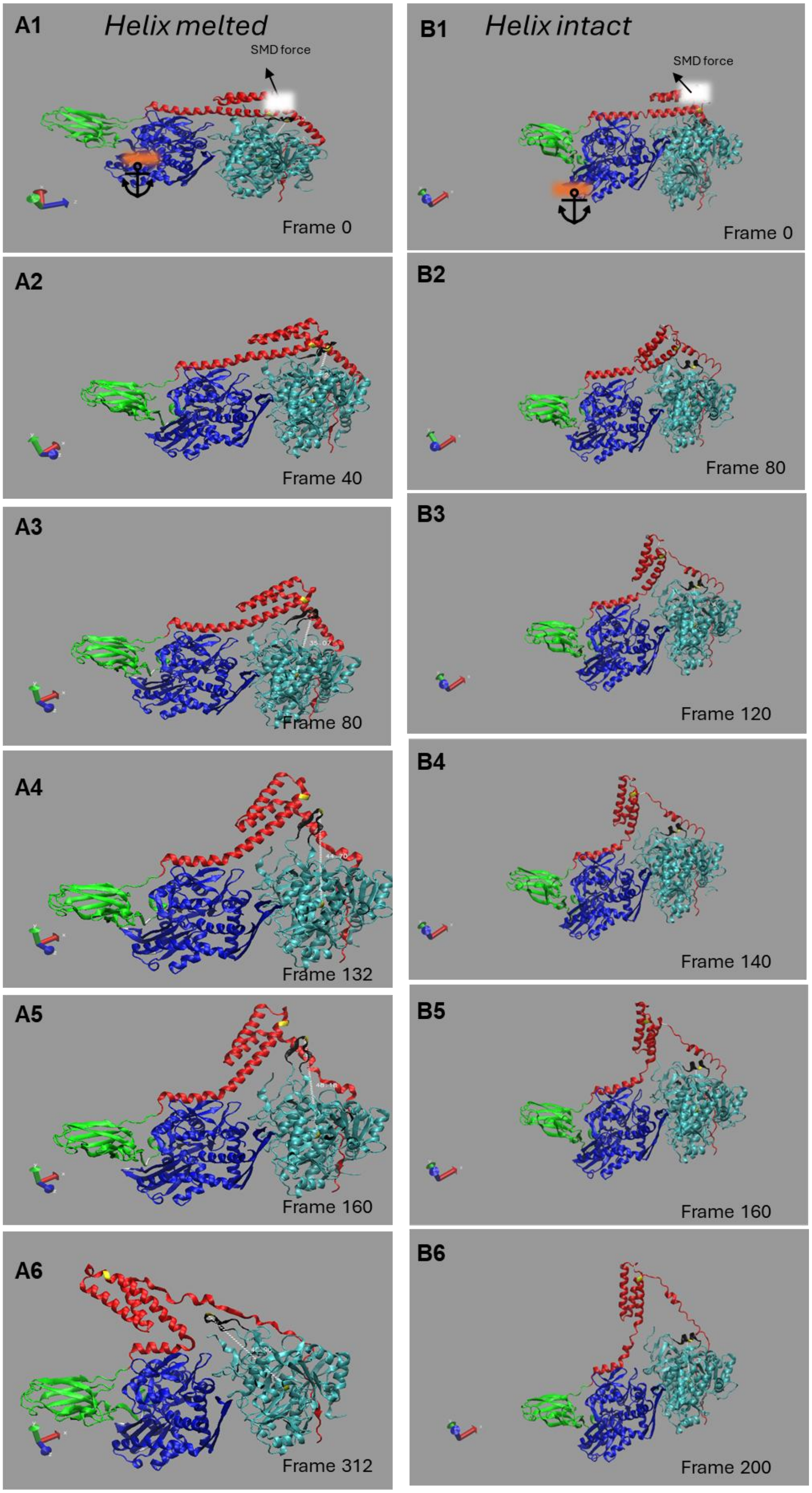

A1
Helix melted
SMD force
Frame 0
B1
Helix intact
SMD force
Frame 0
A2
Frame 40
B2
Frame 80
A3
Frame 80
B3
Frame 120
A4
Frame 132
B4
Frame 140
A5
Frame 160
B5
Frame 160
A6
Frame 312
B6
Frame 200

**Supplementary Figure 8. Model 6.** *Snapshots of the molecular dynamics trajectories illustrate the interaction of the DnaK lid with the Fluc helix 404-411 in its* ***melted*** *state* ***(A)*** *and in its* ***intact*** *state* ***(B****).* The mauve colored-features, labeled with the attached anchor symbol in panels **A1** and **B1** mark a group of residues in the NBD whose Cα atoms were constrained during the SMD pulling of the lid (residues 185, 212, and 388). The white feature on the lid labels the group of residues subjected to the SMD force (residues 555, 560, and 596). The timing for each frame can be determined by multiplying the frame number by 20 ps. Note that following frame 160 (panel **A5)**, the SMD direction changed from (1, 0, 0) to (0.25, 0.15, -1) to better direct the movement of the lid toward SBDβ. The SMD velocity in all simulations involving lid movement, shown in ***A*** and ***B***, was $4x10^{-5}$ Å/step.

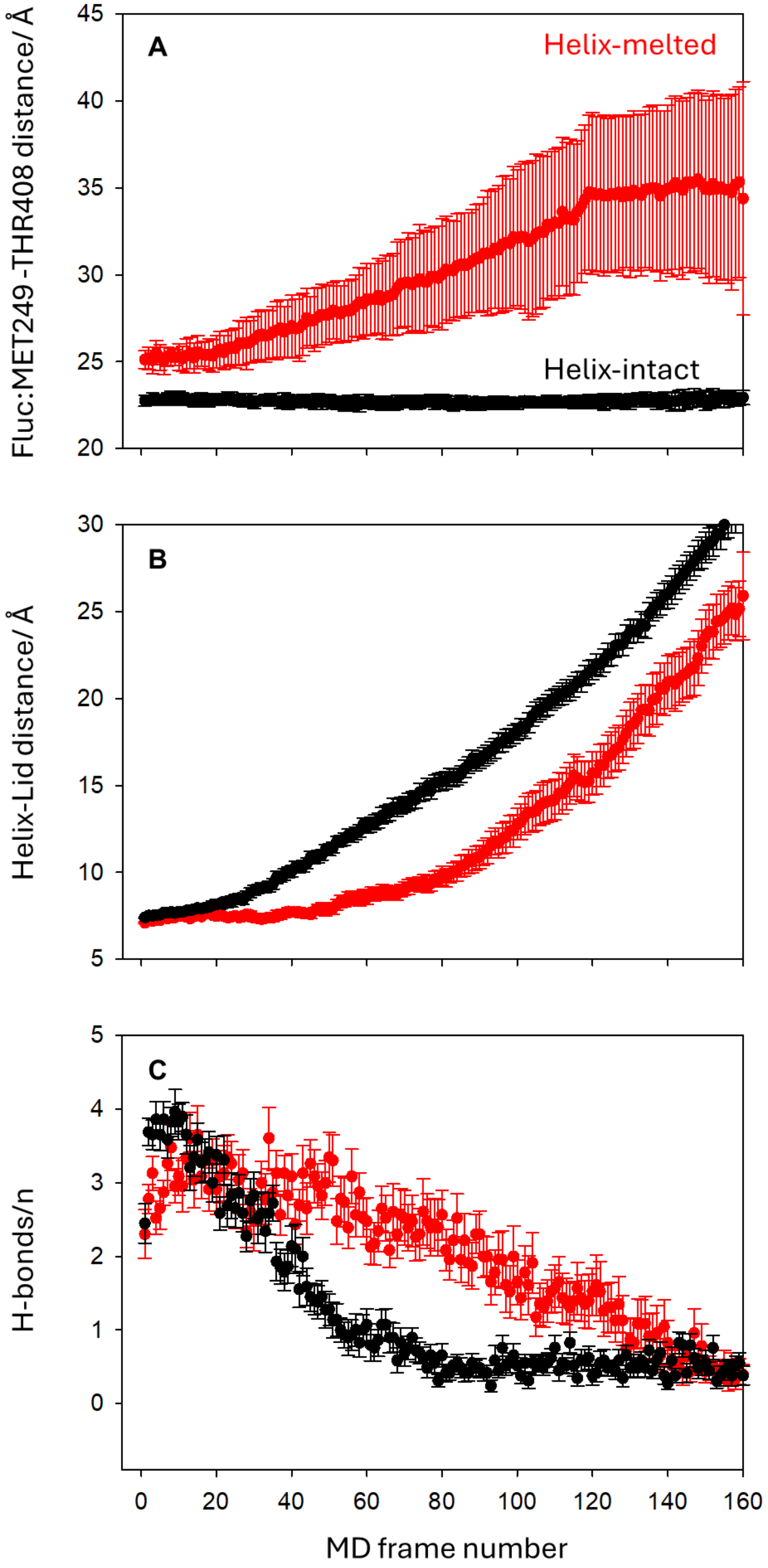

**Supplementary Figure 9. Model 6**. *A comparison of the results of the computational*

*MD experiments capturing the interactions between the DnaK lid and the 405-411 helix of Fluc during lid movement toward SBD$\beta$.* The results for the melted helix (23 independent experiments) are shown in red, and the results for the intact helix (28 independent experiments) are shown in black. The error bars in Fig. 1A depict standard deviations. The error bars in Fig. 2BC mark the standard error of the mean (SEM). In **B**, the distance between the helix and the lid is measured as the distance between the C$\alpha$ atom of Thr408 in the helix and the C$\alpha$ atom of Glu552 in the lid. Hydrogen bonds between Fluc's region encompassing residues 403-413 and Dnak lid's residues 540 to 616 were determined for each MD frame in VMD.
